# Analyses of Eurasian wild and domestic pig genomes reveals long-term gene-flow and selection during domestication

**DOI:** 10.1101/010959

**Authors:** Laurent A.F. Frantz, Joshua G. Schraiber, Ole Madsen, Hendrik-Jan Megens, Alex Cagan, Mirte Bosse, Yogesh Paudel, Richard PMA Crooijmans, Greger Larson, Martien AM Groenen

**Affiliations:** Animal Breeding and Genomics Group, Wageningen University, De Elst 1, Wageningen, WD 6708, The Netherlands; Department of lntegrative Biology, University of California, Berkeley, CA 94720-3140, USA.; Department of Genome Sciences, University of Washington, Seattle, WA 98195-5065, USA; Department of Evolutionary Genetics, Max Planck lnstitute for Evolutionary Anthropology, 04103 Leipzig, Germany.; Durham Evolution and Ancient DNA, Department of Archaeology, Durham University, Durham DH1 3LE, UK.

**Keywords:** Domestication, Gene-flow, ABC, Genomics

## Abstract

Traditionally, the process of domestication is assumed to be initiated by people, involve few individuals and rely on reproductive isolation between wild and domestic forms. However, an emerging zooarcheological consensus depicts animal domestication as a long-term process without reproductive isolation or strong intentional selection. Here, we ask whether pig domestication followed a traditional linear model, or a complex, reticulate model as predicted by zooarcheologists. To do so, we fit models of domestication to whole genome data from over 100 wild and domestic pigs. We found that the assumptions of traditional models, such as reproductive isolation and strong domestication bottlenecks, are incompatible with the genetic data and provide support for the zooarcheological theory of a complex domestication process. In particular, gene-flow from wild to domestic pigs was a ubiquitous feature of the domestication of pigs. In addition, we show that despite gene-flow, the genomes of domestic pigs show strong signatures of selection at loci that affect behaviour and morphology. Specifically, our results are consistent with independent parallel sweeps in two independent domestication areas (China and Anatolia) at loci linked to morphological traits. We argue that recurrent selection for domestic traits likely counteracted the homogenising effect of gene-flow from wild boars and created "islands of domestication" in the genome. Overall, our results suggest that genomic approaches that allow for more complex models of domestication to be embraced should be employed. The results from these studies will have significant ramifications for studies that attempt to infer the origin of domesticated animals.

**Significance Statement:** Though animal domestication has traditionally been viewed as a human-directed process involving small populations of domestic animals and limited interbreeding between wild and domestic forms, but recent zooarcheological insights have questioned this model. By studying domestication in pigs, we demonstrate that complex models of domestication incorporating long-term gene-flow from multiple wild boar populations fit genomic data from modern wild and domestic pigs significantly better than models based on the traditional perspective. In addition, we demonstrate that selection at genes associated with domestic traits countered the effects of the gene flow, thus allowing morphological and behavioural differentiation between wild and domestic populations to be maintained.

## Introduction

The rise of agriculture, which occurred approximately 10,000 years ago, was one of the most important transitions in human history. During the Neolithic revolution, the domestication of plant and animal species led to a major subsistence shift, from hunter-gatherers to sedentary agriculturalists that ultimately resulted in the development of complex societies. The process of animal domestication led to striking morphological and behavioural changes in domesticated organisms compared with their wild progenitors (1). Traditionally, this process has often been viewed as human-directed, involving strong bottlenecks in the domestic population (*i.e.* founder events due to the selection of only a few individuals at the beginning of domestication) and reproductive isolation between wild and domestic forms (2-6). This straightforward model provides an attractive theoretical framework for geneticists, because key events such as the geographic origin and timeframe of domestication are well defined. Thus, the assumption of reproductive isolation eases the interpretation of genetic data from domestic and wild forms. For instance, under this model, geneticists have interpreted phylogenetic affinities of domestic animals with multiple, geographically divergent wild populations as evidence of frequent, independent domestication origins in multiple species (8-13).

However, this view conflicts with zooarcheological evidence that shows that domestication episodes are rare, and that domesticated forms were diffused out from a limited number of core regions (7, 14, 15). Moreover, there is a growing body of empirical and theoretical archaeological work (3, 4, 16) that challenges the simplicity of traditional models. In these new, more complex models, pre-historic domestication of animals is viewed as mainly unintentional (3, 4, 7) and neither reproductive isolation nor strong intentional selection are viewed to be as crucial and widespread as previously thought. Instead, domestication is seen as a long-term, diffuse process (17), involving gene-flow (during as well as post-domestication) between wild and domestic forms (18) and with emphases on multiple, taxon specific, human-animal relationships (3, 4). The possibility of post-domestication gene-flow between domestic animals and their wild progenitors, as well as a lack of strong domestication bottlenecks (18), are key predictions from this novel framework that contrast with more traditional models of domestication. Moreover, extensive gene-flow between wild and domestic forms violates the assumptions of traditional models of domestication and has significant ramifications for studies that attempt to infer the spatial and chronological origin of domestication using genetic data.

Here, we focus on pig domestication using genome-wide datasets of modern domestic pigs and wild boars. Pigs were domesticated independently once in Anatolia (16) and once in the Mekong valley around 9,000 BP (19). Furthermore, ancient mtDNA analyses found that the first domestic pigs in Europe were transported by early farmers from the Levant into Europe around 5,500 BC, concordant with zooarchaeological evidence for a single domestication origin of Western Eurasian domestic pigs (20, 21). However, a few thousand years after their introduction, domestic pigs in Europe had completely lost the Near Eastern mtDNA signatures and instead acquired mtDNA haplotypes typically found in local European wild boars (20, 21). These findings suggest that early domestic populations experienced post-domestication gene-flow from wild boar populations that were not involved in the Anatolian domestication process (7).

Further mtDNA analyses of ancient Anatolian material demonstrated that, by 500 BC, local mtDNA haplotypes were also replaced by haplotypes from European wild boars. This result suggests extensive mobile swineherding throughout Europe and Anatolia (21), consistent with both archaeological and historical evidence, as well as limited management and selection up until the industrial revolution in the 19th century (22, 23). Thus, under a complex model of domestication, mtDNA replacement in ancient European and Anatolian pigs is the result of post-domestication gene-flow, loose pig management and mobile swine herding. We therefore expect such phenomenon to have left a strong signal of gene-flow from wild boars in the genome of modern domestic pigs.

However, while unsupported by any zooarcheological evidence, the observed mtDNA turnovers could also be interpreted as a *de-novo* domestication of a population of European wild boars rather than the result of post-domestication gene-flow from wild boars. Moreover, because of its mode of inheritance and limited resolution, small mtDNA markers provide a very limited impression of gene-flow, making it impossible to test these hypotheses. Thus, the hypothesis of complex domestication in pigs has yet to be tested with the resolution and confidence afforded by unlinked, nuclear markers. In addition, unlike horses and donkeys, intentional interbreeding between pigs and wild boars confers no clear productive advantage and is thought as being mainly unintentional (18). Lastly, there is a clear morphological and behavioural dichotomy between wild and domestic pigs that is evident in modern animals as well as in the zooarcheologic record (24-27). Thus, the possibility of unintentional gene-flow between wild and domestic pigs also raises questions regarding the mechanisms behind the maintenance of traits that differentiate domestic and wild forms.

Here, we fit models of domestication to a genome-wide dataset from over 100 wild and domestic pigs. Our main aim is to ask whether pig domestication follows a traditional, linear model or a complex, reticulate model. More precisely, we assessed whether the zooarcheological evidence for a single, geographically restricted, domestication of (Western) pigs in Anatolia (7, 15, 20) is compatible with the assumption of a traditional model of domestication involving reproductive isolation and strong bottlenecks.

## Results and Discussion

We evaluated the support of multiple models for the domestication of pigs in Europe and Asia. Our analysis focused on 103 genomes from European wild boars (EUW) (8), European commercial / historical domestic (EUD) (Table S1). In addition, this data set comprises multiple populations of Asian wild boar (ASW) and Asian domestic pigs (ASD; Table S1). In order to better understand the early process of domestication, we sampled a range of wild boar populations, from Asia and all major European Pleistocene refugia, rare historical European and Asian breeds, as well as modern commercial breeds. To test key predictions of the complex domestication framework described above, we fit simple but informative models to these genomic data set using Approximate Bayesian Computation (ABC) (see Materials and Methods).

### Testing models of domestication from genome sequences

We first tested the hypothesis of gene-flow between wild and domestic pigs. More precisely, we asked whether reproductive isolation between wild and domestic pigs is compatible with the zooarcheologic evidence that pigs were domesticated only twice, independently in Anatolia and China. To do so, we first evaluated the fit of the traditional model in which domestication is modelled as two parallel events in Asia and Europe. In this model, domestication takes place at time T1 in Europe and T2 in Asia and involves no gene-flow between wild and domestics (reproductive isolation) or between domestics from Asia and Europe (Fig. 1a). We then compared this null model to five other models involving different patterns of continuous mixture: within wild, within domestic, between wild and domestic, etc. (Fig. S1). By comparing these six models (Fig. S1), we found that a model involving gene-flow between domestic and wild (within Asia and Europe) as well as between domestic and domestic (between Europe and Asia) provided a large improvement of fit (Bayes Factor [BF] > 14) when compared to any other model tested in this study (Fig. 1a; Fig. S1). Thus, our explicit modelling framework provides very strong evidence that reproductive isolation between wild and domestics was not maintained during and after domestication in Asia and Europe.

**Figure 1:**
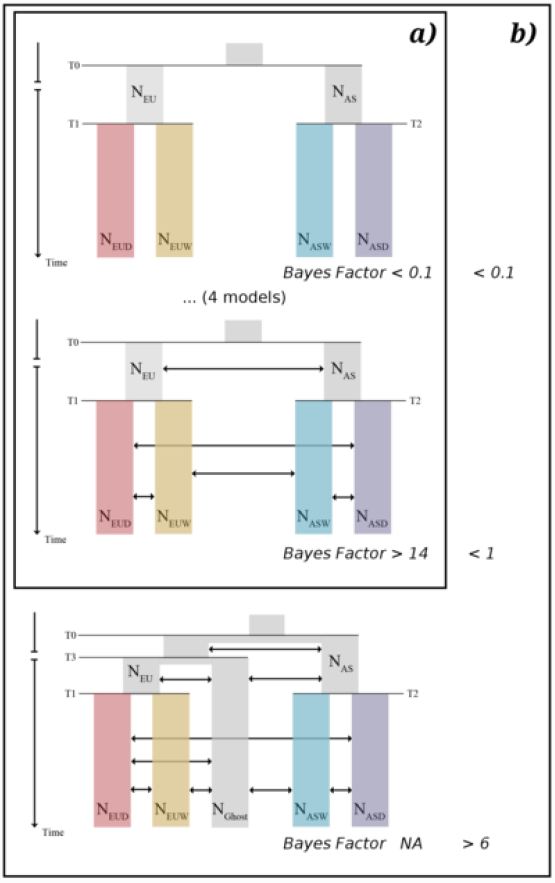
Schematic representation of models tested in this study. Double headed arrows show migrations that were modelled as two independent continuous parameters. **a)** Model testing approach comparing six models. Two models, one without gene-flow (null model; top) and one model with gene-flow between wild (ASW and EUW) and domestic (ASD and EUD) as well as between domestic and wild (full; middle) are displayed. The four additional model tested in this study are displayed in Fig. S1. Bayes Factors in this square were computed without the Ghost model (6 models in total; see b)). **b)** Same as a) but with the Ghost model (bottom). Bayes Factors were computed with all 7 models together.

We further assessed this finding using a dataset of over 600 pigs (from the same populations as in the genome-wide data) that were genotyped on the Porcine SNP60 array (Sl Text). We investigated the historical relationship among these populations using *TreeMix* (28). Our analysis showed that EUD and ASD were both paraphyletic while EUW was monophyletic (Sl Text; Fig. S2). The paraphyly of EUD and ASD is difficult to reconcile with the assumptions of a linear, spatially restricted model of domestication. Instead, this finding provides further evidence of a complex domestication process that involved gene-flow between wild and domestic pigs. Moreover, we found that gene-flow between wild and domestic pigs in Europe was strongly asymmetrical (migration rate from EUW to EUD was much higher than from EUD to EUW; Sl Text; Fig. S3)

Lastly, we also found evidence that Asian and European domestic pigs exchanged genetic material. This is consistent with previous studies and is most likely the result of European importation of Chinese pigs during the 19th century to improve European commercial breeds (23, 29, 30). However, gene-flow between domestic populations (EUD and ASD) was very limited relative to gene-flow between wild and domestics (EUW and EUD; ASW and ASD; Fig. S3). This is not surprising given our sampling strategy that focuses on historical (non-commercial) European domestic breeds that are less likely to be admixed with Asian domestic pigs (23). The small amount of gene-flow between domestic pigs suggests that admixture between domestic pigs had no little influence on our conclusion of gene-flow between wild and domestics (see Sl Text).

Together, these findings demonstrate that domestic pigs do not form a homogeneous genetic group, as expected under a simple human-driven model of domestication. Instead, domestic pigs are a genetic mosaic of different wild boar populations. Thus, modern genetic data from domestic pigs can only be reconciled with zooarcheological evidence for a restricted domestication process if modelled with continuous gene-flow between wild and domestic pigs.

### Demography of pig domestication

We also tested for the presence of a strong population bottleneck associated with domestication. To do so, we estimated the posterior distribution of demographic parameters using 10,000 retained simulations out of 2,000,000. Under the assumption of a simple linear model of domestication with no gene-flow and strong intentional selection by humans, we would expect a strong bottleneck in domestic populations. Overall, we found a population decline in both EUW and EUD (Fig. 2). This is consistent with previous results demonstrating that Pleistocene glaciations resulted in long-term population decline in European wild boars (23, 31-33). However, this population decline was more pronounced in EUW than in EUD (Fig 2). In addition, we found that the effective population size of EUD (Ne-EUD ̴=20,563) was more than twice as large as the effective population size of EUW (Ne-EUW =̴8,497). This is most likely due to a series of strong bottlenecks in the European wild population, caused by over-hunting and loss of suitable habitat (23, 31-33). Together these results do not support the existence of a strong domestication bottleneck in European domestic pigs and instead support the contention that continuous gene-flow from multiple genetically and geographically distinct wild boar populations likely increased the effective population size of EUD.

**Figure 2:**
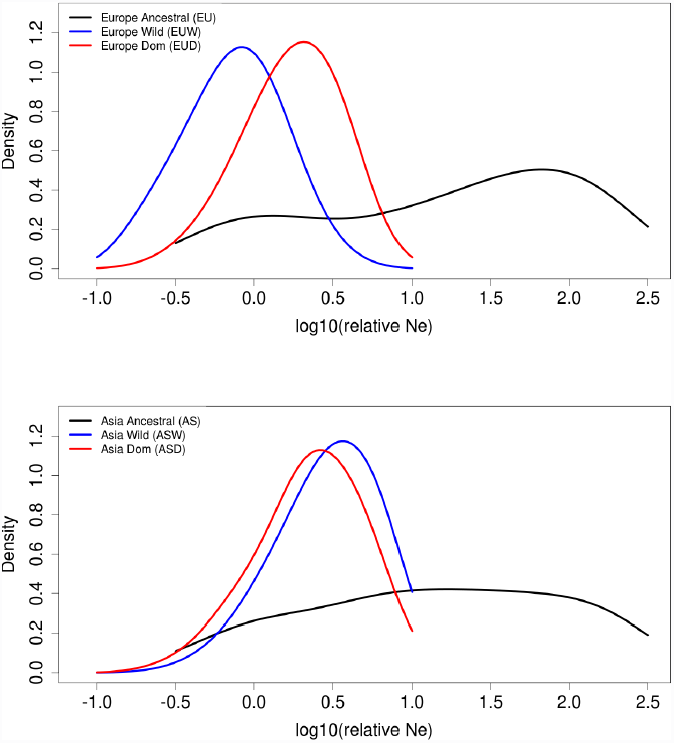
Posterior density distribution of demographic parameters. Population sizes are shown in relative population proportion (the ratio of the current population size over the population size before T0 [Figure 1]).

### Gene-flow from wild boar populations not involved in the domestication process

We showed that a model incorporating continuous gene-flow between wild and domestic pigs is significantly more compatible zooarcheological evidence compared to a traditional hypothesis of reproductive isolation. Despite this fact, we only modelled gene-flow from a population of wild boars that we assumed to be close to the source of domestication. Here, we test the hypothesis that another population of wild boars either extinct (due to hunting pressure and habitat loss (23)) or un-sampled in our analysis (*e.g.* from Central Eurasia, where we did not sample) also contributed to the gene-pool of modern European domestic pigs. To do so, we used a model that is similar to our best fitting model (see above; Fig. 1a) but with an additional "ghost" population that splits from EUW/EUD during the Pleistocene (Fig. 1b) and act as a step between ASW and EUW (migration ASW <-> Ghost <-> EUW but also Ghost <-> EUD; Fig. 1b). This model provided a substantial improvement of fit (BF>6). This result shows that mobile herding of domestic pigs across Europe most likely resulted in gene-flow from a least one wild boar population that was genetically divergent from the population involved in the domestication process in Anatolia.

### Positive selection in domestic pigs

Our analysis shows that gene-flow between wild and domestic forms was a ubiquitous feature of domestication and post-domestication processes in pigs. Thus, extensive gene-flow from wild boars into domestic pigs during and after domestication raises questions regarding the mechanisms behind the maintenance of the clear morphological and behavioural differences observed between domestic and wild pigs. Intentional or unintentional artificial selection could have counteracted the effect of gene-flow and resulted in morphological and behavioural differentiation between wild and domestics.

In order to assess the importance of selection in the genome of domestic pigs in face of gene-flow we conducted a scan for positive selection using *SweeD* (34, 35). *SweeD* computes the composite likelihood ratio (CLR) of a sweep model over a neutral model. Such a test can be very sensitive to demography and migration (36). To correct for effects of demography and migrations we used the 10,000 closest simulations (out of 2,000,000) under our best fitting model in Fig. 1a (without ghost population, see Sl Text) to generate an expected cumulative distribution function (CDF) of neutral CLR and to compute the p-value for all empirical CLR value in the genome (Materials and Methods; Sl Text). We identified 249 and 136 10kbp regions with p<0.01 in the genome of European and Asian domestics, respectively.

First, we examined sweeps private to each population (SI Text). These sweeps in domestic pigs (EUD and ASD) were significantly enriched with GO terms related to multiple developmental processes of bones, teeth and nervous system (Table S1&S2).These terms comprise multiple gene candidates related to height (*PLAGJ*, *NCPAG*, *PENK*, *RPS20* and *LYN* in EUD; Fig. 3a; *LEMD3* and *UPKJ* in ASD) in pigs (37, 38) and cattle (39, 40), nervous system development and maintenance (*NRTN, SEMA3C, PLXNCJ, AAKJ, RAB35, FRS2*) (41-52) as well as genes directly influencing behaviour (*i.e.* aggressiveness and feeding; *APBA2*, *MC4R*, *RCANJ*, *BAJPA3*) (53-60). These results suggest that domestication and/or post-domestication selection for behavioural and morphological traits was important in Asian and European domestic pigs and most likely counteracted the effect of continuous gene-flow in certain parts of the genome.

**Figure 3:**
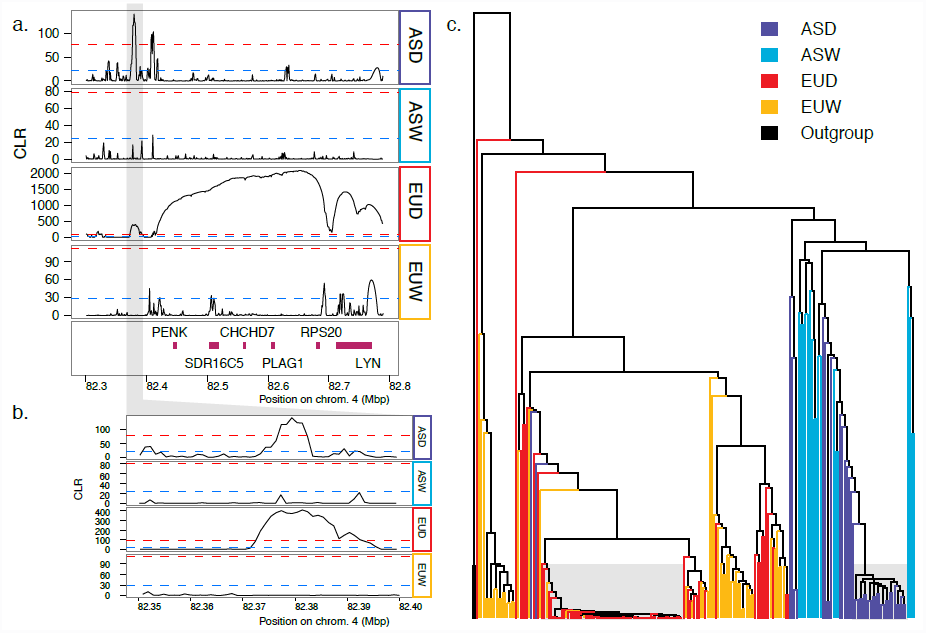
Example of a parallel sweep in ASD and EUD. **a)** Composite likelihood ratio (CLR) values in the PLAG1 region. Dashed blue and red lines represent p=0.05 and 0.01 thresholds respectively. Grey shaded area is the position of the parallel sweep (see b). **b)** CLR values in the parallel sweep region a few thousands base pair upstream of the *PLAGJ* region. **c)** Genealogy of phased haplotypes for the region in Fig. 2b. Shaded area highlights the very short branch lengths that are the result of a sweep. The shaded area on the left side (Europe) contains 64 haplotypes from EUD (>72% of total EUD haplotypes) and 2 haplotypes from EUW (<4% of total EUW haplotypes). The shaded are on the right side (Asia) contains 24 haplotypes from ASD (>54% of total ASD haplotypes) and no ASW haplotype.

However, the mechanism behind this maintenance remains unknown. One possibility is that there was recurrent selection for similar traits. This phenomenon may have resulted in parallel sweeps at the same loci in both ASD and EUD. To investigate this possibility, we looked for signals of parallel selection between the two independent domestication episodes (ASD and EUD). To do so, we identified genes with CLR above the significance threshold in both ASD and EUD but not in ASW and EUW. In order to rule out admixture between ASD and EUD as the cause for observing overlapping significant signal we conducted a phylogenetic analysis in each region separately (Sl Text). The genealogy of some of these regions shows a signal that is consistent with introgression between EUD and ASD (*e.g.* Fig. S7). However, we found one region of particular interest seems to have swept independently in EUD and ASD (Fig. 3). Phylogenetic analysis in this region (Fig. 3b) reveals that ASD and ASW as well as EUD and EUW are mostly monophyletic (Fig. 3c), suggesting an independent sweep in ASD and EUD. Interestingly, while this sweep does not overlap with genes, the region is just a few kbp upstream of the highest CLR in EUD (among others; Fig. 3a). This region has been shown to have a strong effect on body height and stature in pigs (37, 38) and cattle (39, 40). In particular, variation in this region explains up to 18% in body length difference between wild boars and commercial EUD (37). Given the importance of this region for morphology in commercial EUD (38) it is possible that human-mediated selection for similar traits in Asian and European domestic pigs resulted in parallel sweeps at the same loci. Parallel selection of this form may be the responsible for some of the morphological convergence in the two independent domestication events in Europe and Asia. Thus, while the phenotypic effect of this sweep is still unclear, this region provides a particularly interesting candidate to further study the possibility of convergence between ASD and EUD.

## Conclusions

The generation of larger amounts of genomic data with ever-greater resolution is allowing us to embrace the complexity of domestication. The commensurate advancements in theoretical and empirical perspectives is allowing for more sophisticated models to be tested and for a greater understanding of animal domestication. In this study we demonstrated that the assumptions of traditional models, such as reproductive isolation and strong domestication bottlenecks, are incompatible with the zooarcheological evidence of a geographically restricted domestication process in pigs. Instead our model testing approach revealed that continuous gene-flow from wild boars to domestic pigs is necessary to reconcile modern genetic data with the zooarcheological evidence. Moreover, we demonstrated that in Western Eurasia, gene-flow most likely involved at least a second, un-sampled (possibly even extinct) population of wild boars that was quite divergent from the source of domestication. This is most likely the result of mobile domestic swine herding, as predicted by zooarcheologists (18, 21).

Such extensive gene-flow from wild boars raises questions regarding the maintenance of morphological and behavioural traits in domestic pigs. Our study reveals extensive evidence of selection at candidate genes that influence anatomical and nervous system development, suggesting that selection may have counteracted the homogenizing effect of gene-flow and maintained the genetic basis for the morphological and behavioural dichotomy observed between wild and domestic pigs. In addition, our results show that regions close to genes governing morphological traits have been subject to selection in parallel in Asia and Europe. In the context of speciation, studies focusing on systems in which gene-flow is common have identified genomic regions that show excessive inter-specific divergence (*e.g..* 61). These studies have suggested that such regions may be resistant to gene-flow and likely allowed for the maintenance of species characteristics (genomic “islands of speciation”) (61). Here we hypothesise that an analogous process is taking place during pig domestication. By recurrently selection for similar traits through artificial selection, but allowing for gene-flow, farmers have created genomic “island of domestication”, that we define as genomic regions governing domestic traits and thus less affected by gene-flow from wild boars. However, it is unclear whether these sweeps involved recurrent selection of different haplotypes from standing genetic variation in wild boars or are the result of selection from *de-novo* mutations. Thus, our results highlight a list of candidate genes that will provide further studies with the means to further test these hypotheses.

Lastly, it is important to underline the limitations of modern DNA and traditional domestication models to determine the origin and time of domestication of animals, as well as to identify the genes involved during domestication. Indeed, extensive gene-flow clearly violates the assumption of traditional models and likely eroded most of the signal to infer time and geographic parameters (62-64). It is therefore important to apply caution when conducting comparative analyses of modern genetic material from wild and domestic animals. However, future sequencing of ancient DNA, together with more realistic modelling frameworks, such as the one presented here, will provide the necessary information not only to determine the origin and time of domestication of animals but also to identify genes involved during domestication and will ultimately significantly enhance our knowledge of this fascinating evolutionary process.

## Materials and Methods

### Sample collection and DNA preparation

Blood samples were collected from a total of 622 individuals, 403 European domestics, 92 Asian domestics, 103 European wild boars and 23 Asian wild boars and a Javan Warty pig (*S. verrucosus*), used as an outgroup (33). For full description of the samples see Table S1. DNA was extracted from the blood samples with QIAamp DNA blood spin kits (Qiagen Sciences). Quality and quantity of DNA extraction was checked on a Qubit 2.0 fluorometer (Invitrogen). Single nucleotide polymorphism (SNP) genotyping was performed with the Illumina Porcine 60K iSelect Beadchip. For the genome resequencing, we used 1-3 ug of genomic DNA to construct libraries (insert size range 300-500 bp). Library preparation was conducted according to the Illumina library preparation protocol (Illumina Inc.). Sequencing was done on Illumina Hi-Seq with 100 and 150 paired-end sequencing kits.

### Alignment and variant calling

All samples selected for genome sequencing were sequenced to approximately 10x coverage (Table S1). Reads were trimmed for a minimum phred quality > 20 over three consecutive base pairs and discarded if shorter than 45 base pairs. Alignment was performed with Mosaik Aligner (V. 1.1.0017) with the unique alignment option to the Porcine reference genome build 10.2. Variants were called using GATK Unified Genotyper version 2.8 (65). We used a prior of 0.01 for the probability of heterozygous calls (32).

### Approximate Bayesian Computation

104 genomes were used for the ABC analysis. Simulations were performed on 100 10kbp unlinked loci. Backward coalescent simulations with recombination were performed using *ms* (66) under 7 models (Fig. S1). For model testing purposes, we ran 200,000 simulations per model. Summary statistics were computed on observed and simulated data using using libsequence (67). We compared all models simultaneously (68) using a standard ABC-GLM approach as implemented in *ABCtoolbox* (69). For parameters inference we ran 2,000,000 simulations under the best fitting model. We extracted 10 Partial Least Square (PLS) components from the 93 summary statistics in the observed and simulated data (70). We retained a total of 10,000 simulations closest to the observed data and applied a standard ABC-GLM (71).

### Exploratory analysis using SNP array

We used *TreeMix* (28) to build a maximum likelihood (ML) population tree from the 60K SNP dataset. We generated 10 replicates (with different seeds) and selected the run with the highest likelihood score. The PCA analysis was performed using *flashpca* (72).

### Selection scan

We used SweeD to detect sweeps (35). To obtain critical threshold values (p-values), we used a posterior predictive simulation (PPS) approach. We simulated 2 replicates of 3Mbp each using the parameters of the 10,000 closest retained simulations from our ABC analysis (20,000 simulations). Simulations were run using macs (73). We derived a critical threshold for observed CLR in each population using the cumulative descriptive function (CDF) derived from the CLR distribution that was obtained from the PPS. All regions with p<0.01 were selected for further analysis.

## Acknowledgments

We thank Ben Peter for his help and guidance during the model-fitting step of the analysis as well as for kindly sharing his code. We are also indebted to Daniel Wegmann for providing us with the latest version of *ABCtoolbox*. We also thank Konrad Lohse for his insights during the conception of the project. This project is financially supported by the European Research Council under the European Community’s 256 Seventh Framework Programme (FP7/2007-2013)/ERC Grant agreement no 249894. JGS was supported by National Institutes of Health grant R01-40282 and National Science Foundation postdoctoral fellowship DBI-1402120.

